# Evolving better RNAfold C source code

**DOI:** 10.1101/201640

**Authors:** W. B. Langdon

## Abstract

Grow and graft genetic programming (GGGP) can automatically evolve an existing state-of-the art program to give more accurate predictions of the secondary structures adapted by RNA molecules using their base sequence alone. That is, genetic improvement (GI) can make functional as well as non-functional source code changes.

## 1 Introduction

Biology is now a data rich science. The central dogma of Biology states that the fundamental information for all forms of life is transcribed from DNA into messenger RNA, which intern is translated into protein. Like DNA, RNA is a long chain polymer principally composed of 4 bases (G, U, A, and C). Each RNA molecule’s sequence of bases is known as its primary structure. Much of the interesting Biology occurs when RNA is a single strand (unlike the more stable double stranded DNA helix). Like DNA the four bases can form relatively weak temporary bonds with their complementary base. (C pairs with G, and A with U.) In some commercially important species, perhaps half the DNA is transcribed into RNA but only a tiny fraction of that is translated into protein. Other than conveying information for protein manufacture, there are some well known biological uses of RNA. For example, some RNA molecules are known to act as enzymes which catalyse reactions between biomolecules. Also it is thought some of the transcribed RNA is used to regulate gene expression. Much of the chemistry of biomolecules is determined by their three dimensional shape. These areas are relatively new, and this, and other uses of RNA, have sparked renewed interest in RNA and its structure.

**Figure 1:**
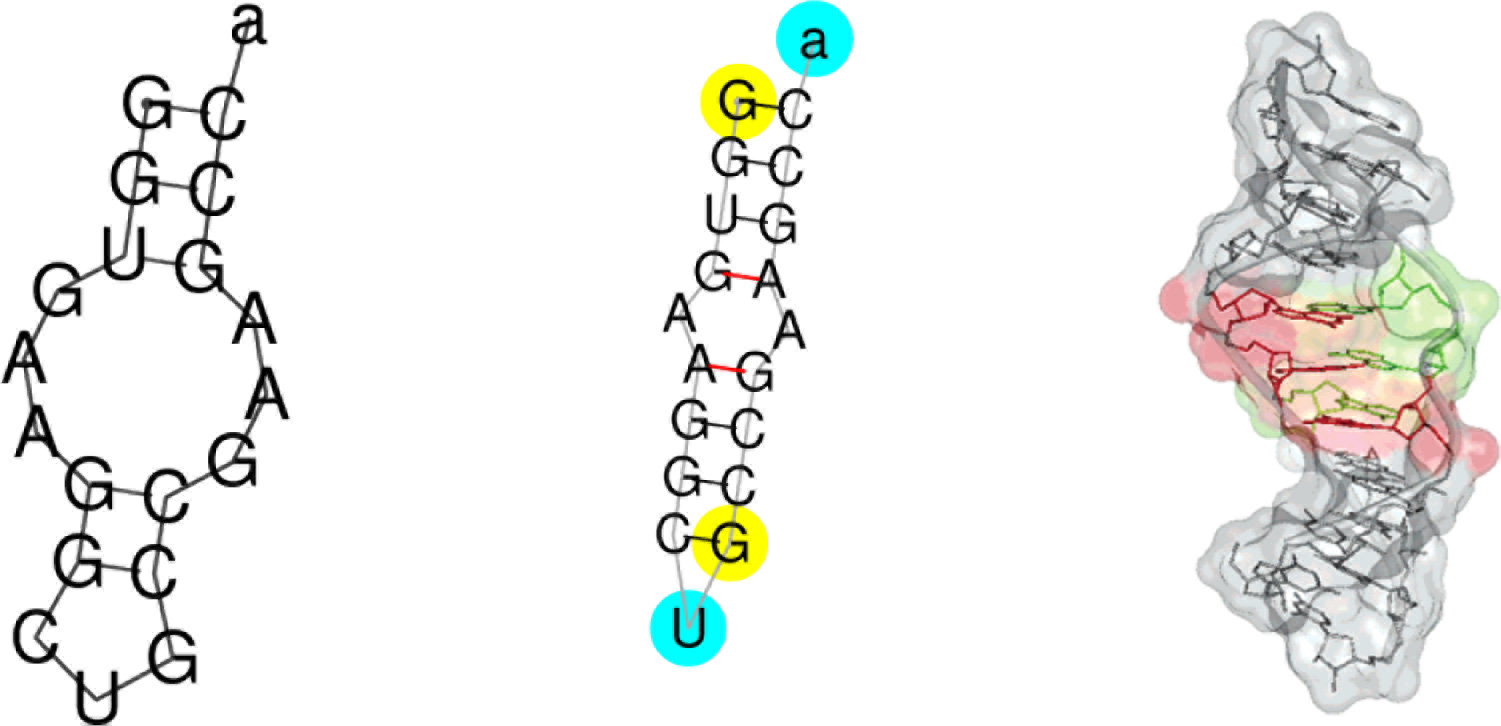
Folding pattern (secondary structure) for RNA molecule PDB_00996 (sequence GGUGAAG-GCUGCCGAAGCCa). Left: prediction made by mutated RNAfold (code changes given in Section 3.3).This structure has a Matthew’s Correlation Coefficient (MCC, page 4) of 0.784133 with the true structure (centre) as recorded by RNA_STRAND v2.0 [1]. (See also Figure 5, page 7.) Red lines in the middle image indicate non-standard RNA base-to-base binding, which RNAfold assumes does not happen [2]. Right: one of true three dimensional structures [3].

First principles computer programs have had some success at predicting RNA secondary structure, i.e., the folding patterns of real RNA molecules (see Figure 1). Mostly these are based on estimating the energy associated with each possible secondary structure using dynamic programming and assuming the molecule will adapt the structure with the lowest energy. In principle considering all possible RNA folding patterns is exponential but many patterns can be discarded as not being biologically plausible. For example, the structure of many RNA molecules is known and very few known structures have knots. Indeed in RNA molecules of known structure, on average 95% of the structure is also free of pseudoknots [1, Table 1]. It is common for structure prediction software to assume that RNA contains no knots [4]. Such dynamic programming based approaches scale approximately O(*n*^3^), where *n* is the number of bases in the RNA molecule. Nonetheless great savings can be made by running such algorithms on parallel hardware [5]. The state of the art in RNA secondary structure prediction is RNAfold from the ViennaRNA package [6]. It has been widely used, for example it is a key component of the eteRNA citizen science project [7].

Grow and graft genetic programming (GGGP) [8, 9, 10, 11, 5, 12, 13] has built on existing genetic improvement (GI) work [14, 15]. GI has been used to improve the performance of existing software, e.g. by reducing runtime [16], energy [17] or memory consumption [18], but (apart from software transplanting [19] and automatic bug fixing [20]) it usually tries not to change programs’ output. GGGP builds on GI which is very much hands off (i.e. let the computer solve the problem) by admitting more human assistance. We applied GGGP to RNAfold [21]. Using traditional manual methods to identify performance critical components, recoding them using Intel’s SSE vector instructions and then using computational evolution to further improve the new code. Unfortunately, evolution only found small increments on the human code and so, so far, only the manually written code has been adopted. However it has been included into the standard ViennaRNA package since 2.3.5 (14 April 2017)^1^. It is also being used by the eteRNA development team internally [22].

The next stage was to apply GGGP to improve the accuracy of RNAfold’s predictions. Notice here we allow (nay encourage, require) evolution to change the output of the program. I.e. to make functional changes, The next section describes our GP [23] system, whilst Section 3 describes the results of applying it to RNAfold’s C source code. We conclude (Section 3.4) that instead of manipulating the code itself, perhaps we should apply evolution directly to the constants [18] used to specify the energy of each potential way RNA might fold up.

## 2 Genetic Improvement System

Evolution uses a BNF grammar to control the mutations it can make. The grammar ensures that after each mutation the new code is still syntactically correct [16, 24, 25, 13, 26]. The grammar was created automatically line by line from the 74 RNAfold .c source files (release 2.3.0, 60 636 lines of code). To simplify fitness testing, the file holding the `main(), RNAfold.c`, is excluded from the BNF grammar in order to ensure evolution cannot modify it.

### 2.1 Profiling with GNU gcov to Direct Evolution to Used Code

The goal of preprocessing is essentially to direct evolution to mutate only code that is in use. Whereas in [16] we used an especially instrumented version of the automatically generated grammar to record which lines of source code where used most heavily, here (like [21]) we used the GNU gcov profiling tool.

We compiled the unmodified released sources using the GNU C compiler (v4.8.5) with command line options `-g −pg −ftest-coverage` and `-fprofile-arcs` and ran the resulting `exe` on all 1555 training RNA sequences (using the same command line as will be used in fitness evaluation, section 2.7, i.e. the defaults plus `––noPS`). Then the profiler was run (`gcov *.gcda`) and the line numbers of the lines used were extracted from `*.gcov` using gawk. The line numbers are used to set binary (0 or 1) weights as to the importance of which rules to mutate when the BNF grammar is used by evolution.

### 2.2 BNF Grammar Types

Each line of the source code is represented by one or more rules in the BNF grammar. Evolution can change the contents of the code but it cannot change its structure. So the .h and .inc include files are excluded, and function and variable declarations cannot be changed. However, evolution can substantially change the code inside functions and between flow control statements including whether functions are called or not. Each variable rule belongs to one of seven types. Mutations only move code between rules of the same type.

The BNF grammar contains 94 810 rules, of these in principle 34154 can be modified. Most (19 576) are lines of C source code which can in principle be moved, inserted or substituted for other lines of source code within the same source file. (There are 74 .c files.) There are 6 375 IF rules (whose conditions can be replaced by the conditions from other rules within the same source code). 2603 `for` loops whose three components can be substituted by the similar part of other `for` loops. 324 WHILE rules and 70 ELSE rules. However these numbers fall dramatically when we include the weights (see Section 2.1). Using the binary weights imposes the restriction that a line of code must have been used at least once when processing all 1555 examples in the RNA_STRAND training data in order to be a target for mutation. (Unused lines, within the same source file, can still be inserted before or used to replace target lines.) Only 20 .c files are used when processing RNA_STRAND. The number of simple source lines that can be modified or used to modify others falls to 4518. There are corresponding reductions in the numbers of rules of each type: 1904 IF, 371 `for`, 27 WHILE and 10 ELSE.

We force loop termination via the Unix `limit cputime` command (see also page 4).

### 2.3 Sandbox Protection Against Rogue Mutants

As well as ensuring evolution cannot cause processes to continue indefinitely we checked the source code for system calls.

RNAfold makes limited use of files, so it was straightforward to confirm that the grammar does not allow evolution to modify how files are used. Especially it cannot change which files are used or how they are opened (although of course it can change the contents of files that RNAfold normally writes to).

In principle, evolution could still cause problems by causing an existing RNAfold output file to grow enormously, potentially filling the disk. However such a rogue mutant would still be limited by the CPU limit. Disk quotas can also be used. We had intended to run the fitness evaluation on a scratch disk to isolate it however the problem never arose in practise and GGGP was run on a conventional shared network drive (albeit with disk quotas enabled).

### 2.4 Avoiding Phenotypic Duplicates – Ensuring a Change is made

For simplicity (unlike some of our work with parallel GPU code using CUDA, e.g. [11]), we did not enforce each new individual to be phenotypically unique. However to ensure mutation has had some impact before fitness testing we did use a simple bit wise binary comparison of the mutated exe to ensure they were all different from the unmutated program. In the run to be described in Section 3, only 16 mutant programs were identical. It seems reasonable that the unmutated exe is the most commonly binary duplicated phenotype, suggesting there is little to be gain from enforcing exact binary phenotype diversity. Indeed since only 16 duplicates were detected, this check could probably be remove with little loss.

### 2.5 Initial Population

The initial population of 500 × 10 GP individuals is created using mutation. The mutation script ensures each member of the initial population is unique.

#### 2.5.1 Over Sized Population

For simplicity (unlike some GGGP work on CUDA code [11]) we do not use scoping rules. Therefore, there was a fear that many mutants would fail to compile. Since the expensive part of fitness evaluation is running RNAfold and compiling code which contains errors is relatively cheap, it was decided to create ten times as many individuals as needed at the start of each generation. This was done using the usual mutation and crossover operations (see next section). However, on average (including the initial generation) only 32% of mutants failed to compile (see also Figure 3).

### 2.6 Mutation and Crossover

The genetic operators used are those we used in [11]. Briefly, each member of the breeding pool, i.e. the top 250 individuals from the previous generation, is allocated two children. One child is created by mutating the parent. The second is created by crossover between it and another parent randomly selected from the breeding pool.

New mutations are appended onto the end of current mutation list. There are three types of code mutation: 1) delete a line, 2) copy and replace the target line and 3) copy and insert a line before the target line.

Two point crossover (similar to Koza’s sub-tree crossover [27]) creates a new child from the lists of code changes from its two parents. If crossover is unable to create a new unique offspring after a five tries, the child is created by mutating the first parent instead.

### 2.7 Fitness Function

Every generation, in order to be able to compare the genetically improved code with the original, the first program compiled and run is always a copy of the default (manually written) code. I.e. at least 501 programs are compiled and run (see also Section 2.5.1.

When a compilation fails, or the resulting program exe is identical, the mutant is not tested and cannot be a parent for children in the next generation.

The mutant exe is run on all the training data. I.e., one 1/3^rd^ of RNA_STRAND which are less than 155 bases long. This means running the mutant’s exe up to 681 times. In retrospect this is perhaps too many. For example, in [16] we used just five but these were randomly changed every generation.

As the unmutated RNAfold is fast and all the of RNA molecules used for training are small, the per fitness case CPU limit was the same in all cases (1 second).

RNAfold was run with option `––noPS` to suppress the production of nice pictures of the predicted structure. (The defaults were used for all other options.)

RNAfold produces its prediction as a text string composed on nested brackets (to indicate pairs of bases which bind together) and “.” (unbound bases). This was piped in the standard Vienna-RNA (2.3.0) utility `b2ct` which converted the bracket string into a .ct file format. The output from `b2ct` was piped into a comparison gawk script which calculates the Matthew’s correlation coefficient 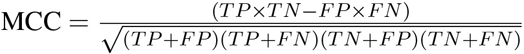. Where:

- *TP* = true positives, number of predicted pairs which are in RNA_STRAND’s.ct file.
- *TN* = true negatives, total number of possible pairings not in *TP*, *FP* or *FN*. I.e. *TN* = *n*(*n* − 1)/2 – *TP* – *FP* – *FN* (where *n* is the length of the RNA molecule).
- *FP* = false positives, number of predicted pairs which are not in RNA_STRAND’s.ct file.
- *FN* = false negatives, number of pairs in RNA_STRAND’s.ct file but not in the mutant’s prediction.

Naturally *TN* tends to be large, hence we follow [6] and use Matthew’s correlation coefficient as it deals well with large class balances [6].

Should the mutated program fail to create a valid prediction on any training case, fitness testing of it is aborted. Missing or invalid predictions are given an MCC of −1.

The population is sorted by the number of valid MCC each mutant has and then by their mean MCC (across all 681 training cases). The top half of the list are then selected to be parents of the next generation. subject to: they must have produced at least one prediction which has an MCC. In all generations, see next section, the number of mutants eligible to be parents exceeded half the population (mean 411.4). Similarly, they all produced a valid MCC on all the test cases. I.e. a mutation which did not produce a valid prediction on all 681 test cases was never selected to be a parent.

**Table 1:** GGGP to improve RNAfold’s secondary structure predictions by mutating its C code sources.

|  |  |
| --- | --- |
| Representation: | Variable list of replacements, deletions and insertions into BNF grammar |
| Fitness: | Compile modified code. Run on training data from RNA_STRAND with less than 155 bases (681 molecules). Compare its answers with the real structure using MCC. See Section 2.7. |
| Population: | Panmictic, non-elitist, generational. 500 members. (Create ten times as many individuals with mutation and crossover, mutation only in initial generation, as needed but only accept the first 500 which compile. See Figure 3.) |
| Parameters: | Initial population of random single mutants. 50% truncation selection. 50% two point crossover, 50% mutation. No size limit. |

**Figure 2:**
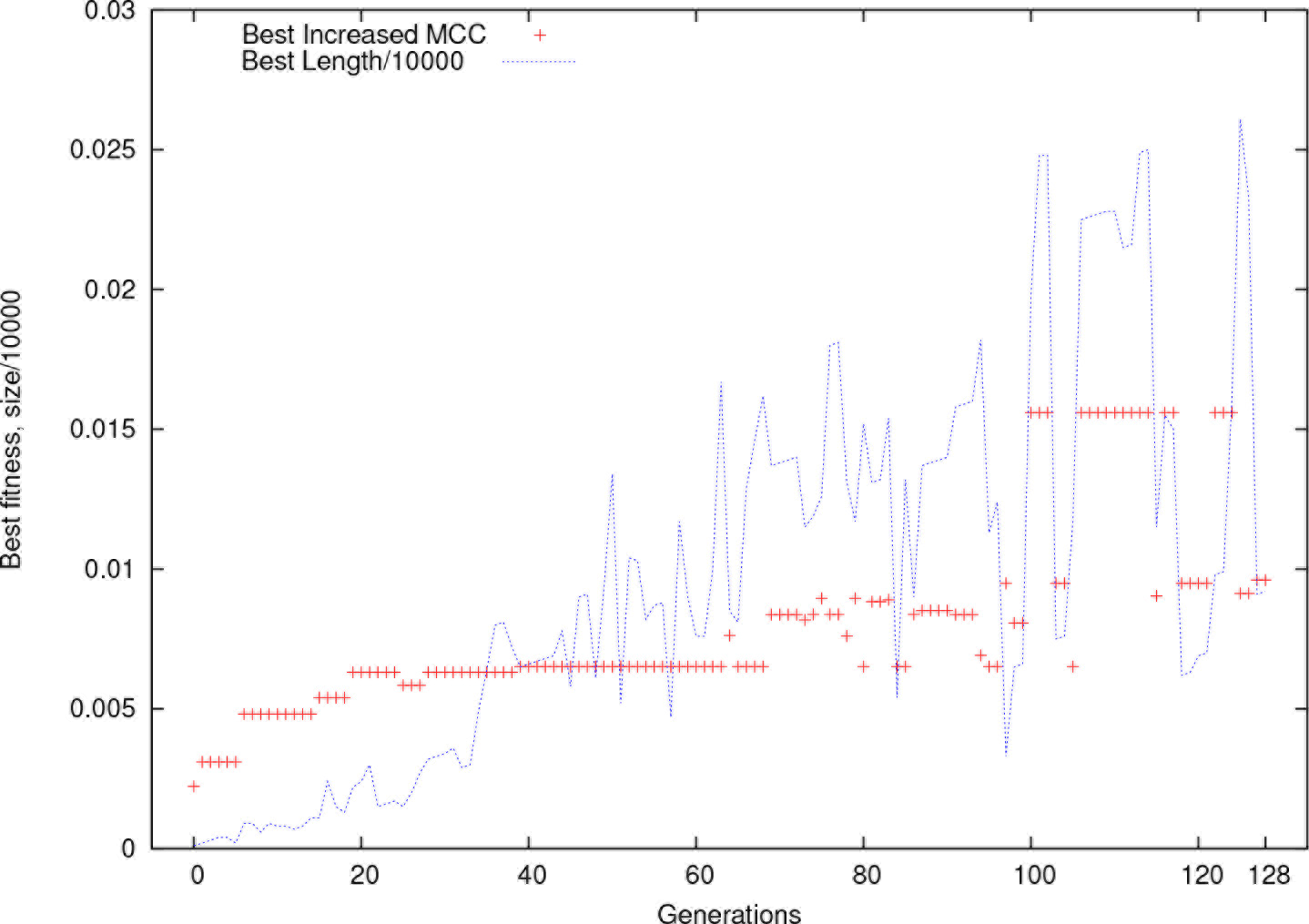
Best in generation for GGGP run (population 500). Crosses + show increase in MCC averaged across 681 training examples. (I.e. increase above released code performance 0.663946). Dotted line shows size of best in population (note change of scale). Run stopped after generation 128, when best individual made 92 code changes.

## 3 Results

The GGGP was run with a population of 500 (see Table 1). The run was aborted after 22 days when the 3.6GHz Centos 7 desktop was rebooted. The evolution of training performance is given in Figure 2.

### 3.1 Post Evolution Tidy

It is common practise in GI to allow evolution to progressively increase the number of mutations per individual [28]. Indeed, as it was anticipated that many changes would be required in large programs, in some cases such bloat was deliberately encouraged [16]. (In fact it turns out in many cases that only a few changes are actually required to make spectacular improvements [16].) Therefore it is common to include a simple post evolution phase where each mutation is tested one at a time to see if it can be removed without loosing the benefit of the whole change [16].

The number of possible combinations grows exponentially with the length of the evolved individually, therefore we use a simple hill climbing scheme [28]. Start at the front of the best evolved individual in the population and progressively remove each mutation and test the new individuals on the training set. If it fails or it performs worse than the evolved individual, the deleted mutation is restored, otherwise it is deleted permanently. Continue until the end of the evolved individual so that each part has been checked.

**Figure 3:**
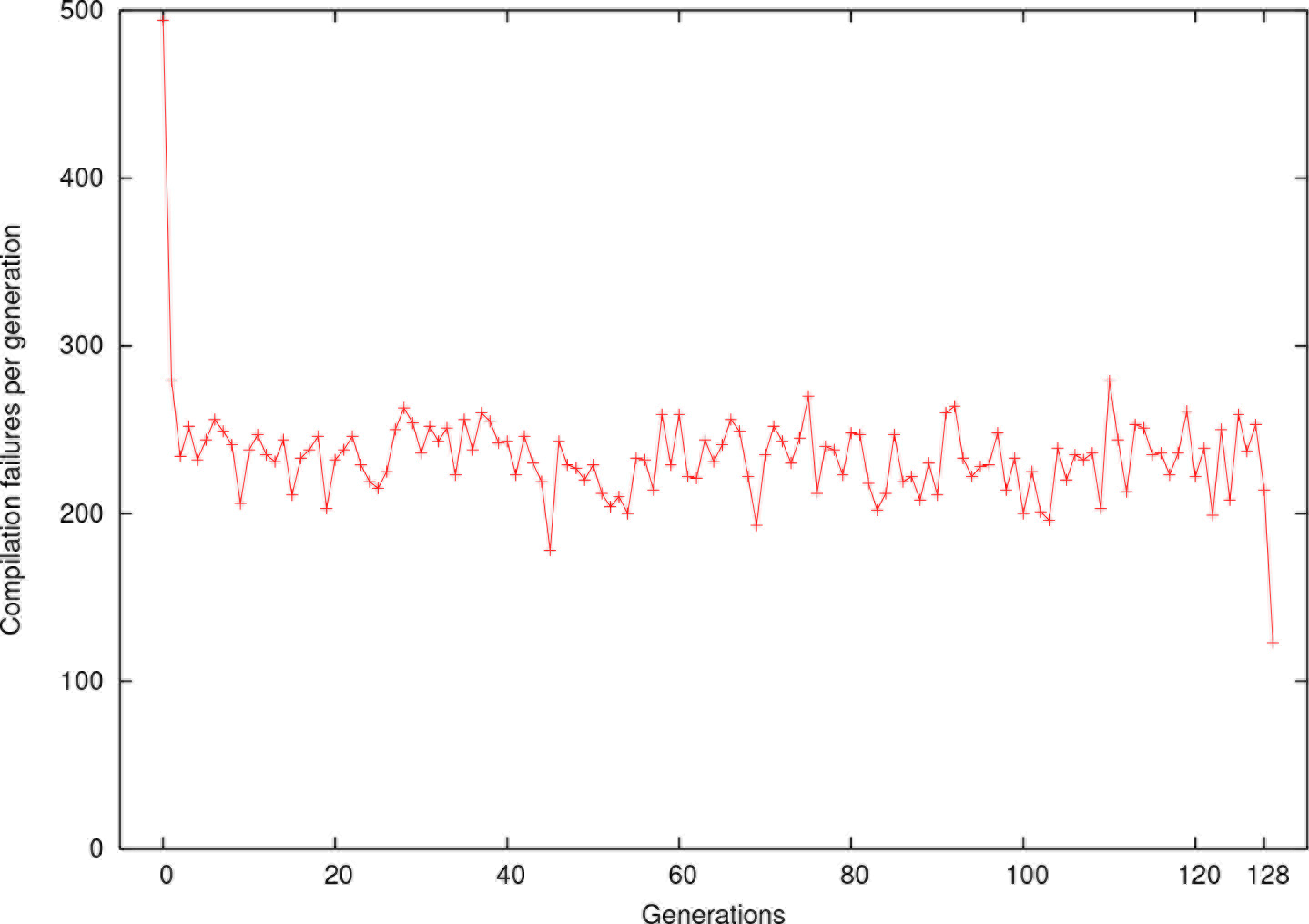
Apart from the initial generation, approximately two thirds of GGGP RNAfold mutants compile.

Another strategy is to start with an empty individual and progressively add mutations to it, retaining only those that improve performance [24, 25].

By generation 128 the best in the populations scored 0.673551 on the 681 short RNA molecules used for training but had grown to 92 code changes. Running our simple hill climbing procedure removed 88, leaving just 4 mutation. The cleaned up individual is `<for1_params.c_291><for1_params.c_277> <_params.c_350> <_params.c_339> <_model.c_860>`

### 3.2 Generalisation Performance

The cleaned up individual retains its slight performance edge on the original code even when used to predict the shapes of RNA molecules from RNA_STRAND not used in training, including molecules of all lengths, see Figure 4.

### 3.3 Code changes

#### 3.3.1 <for1_params.c_291><for1_params.c_277>

This mutation replaces the first part of the `for` loop on line 291 with that from the `for` loop on line 277 in file `params.c.` (New code `for`(`i = VRNA_GQUAD_MIN_STACK_SIZE;i<=MAXLOOP;i++`).) This causes the loop control variable `i` to be initialised with `VRNA_GQUAD_MIN_STACK_SIZE` (2) rather than not being initialised but instead being set to the value which caused the previous loop to terminate 30. Thus the mutation causes `params->bulge[2..30]` and `params->internal_loop[2..30`] to be recalculated overwriting their existing values, which were derived from values stored in arrays `bulgedH[i]` and `internal_loopdH[i]` respectively.

**Figure 4:**
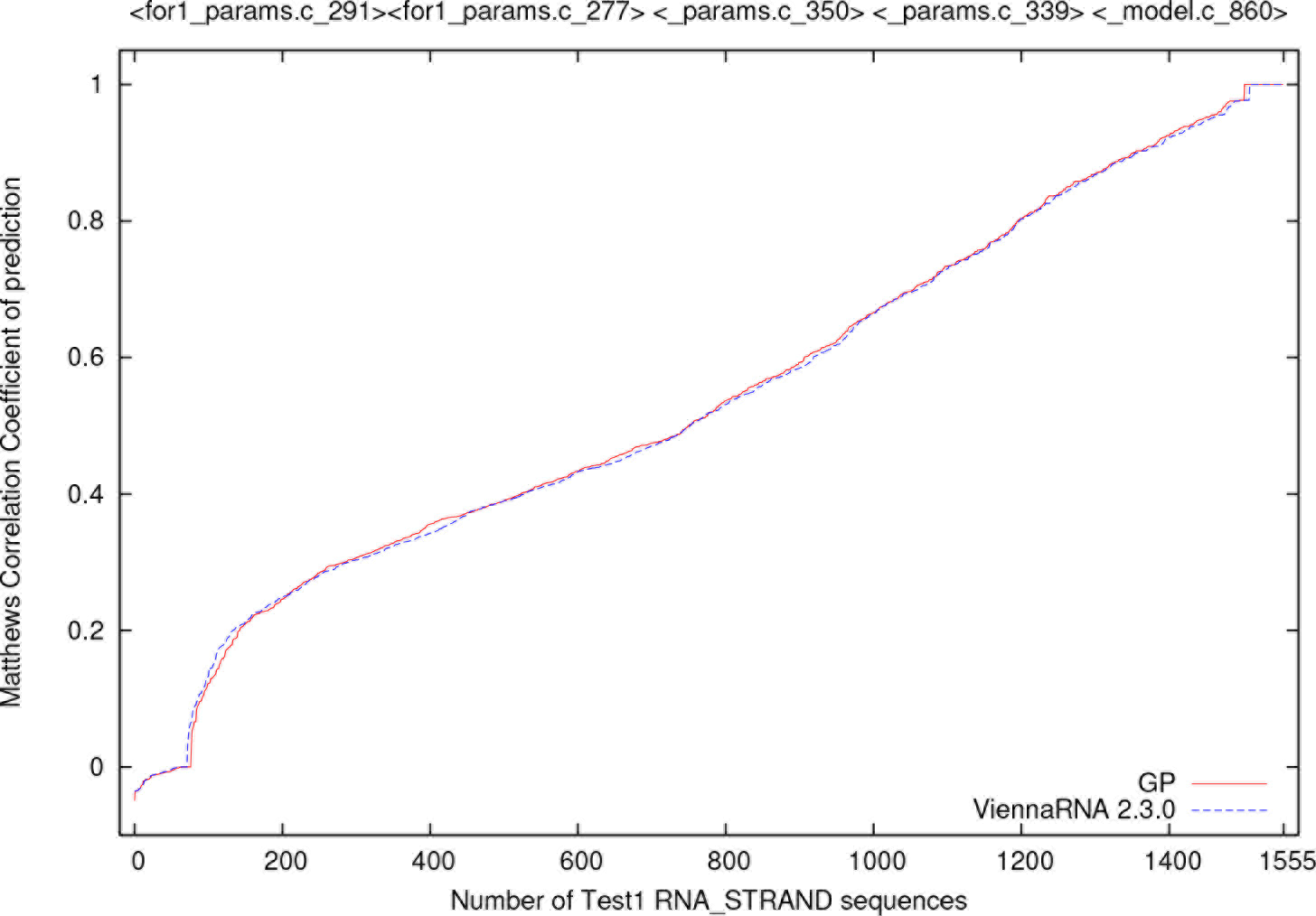
Performance of evolved (best gen 128) version RNAfold (solid line) after automatic clean up Section 3.1 on 1555 RNA_STRAND molecules not used in training. It does significantly better (*p* = 0.00008612 non-parametric two tailed sign test) than the unmutated code (dashed line). Figure 5 plots data as X-Y scatter plot.

**Figure 5:**
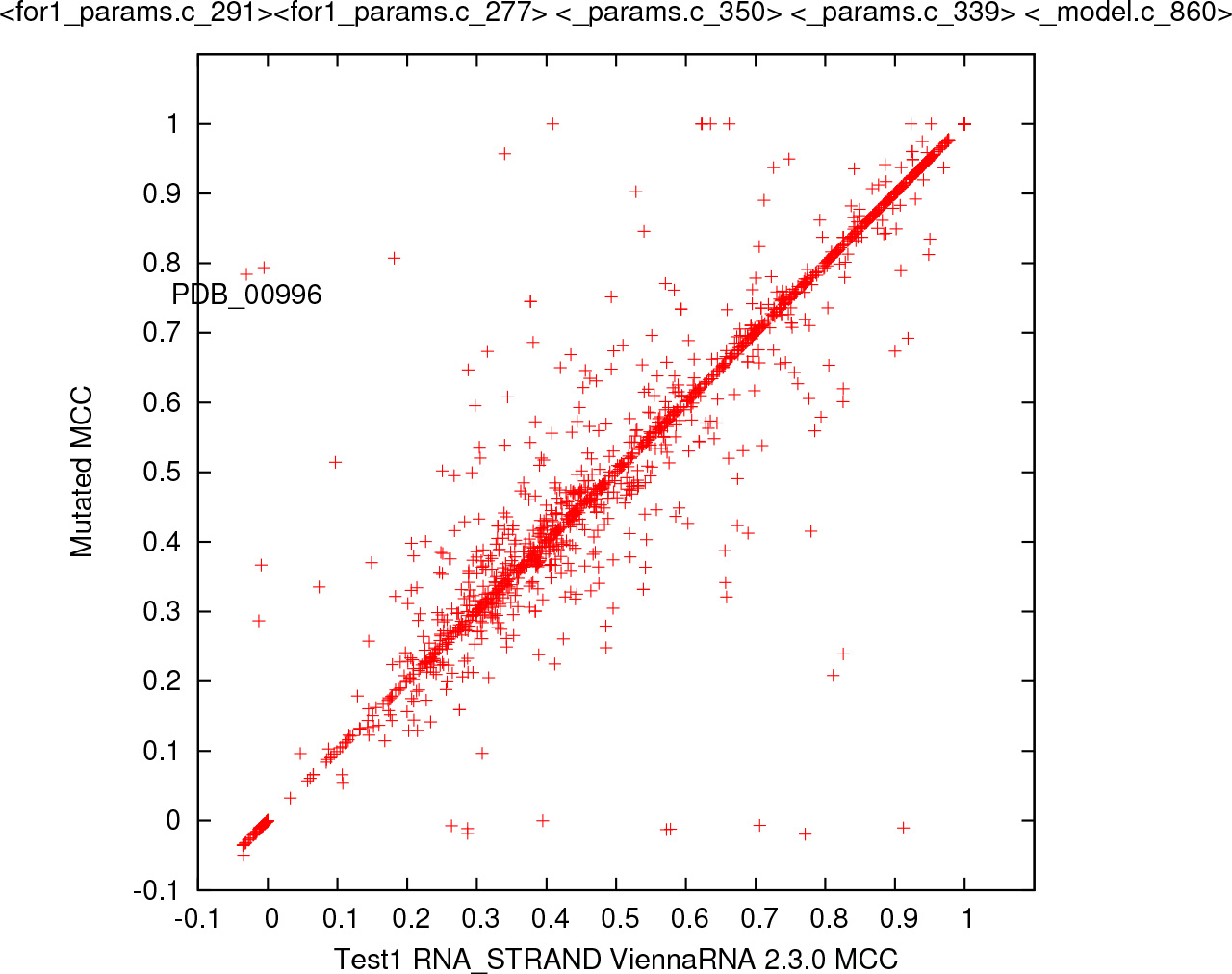
Performance of evolved RNAfold (vertical) v. unmutated code (horizontal). Data as Figure 4.

#### 3.3.2 <_params.c_350>

This causes line 350 in `params.c` tobedeleted: `dd = dangle5_dH[i][j] – (dangle5_dH[i][j] – dangle5_37[i][j])*tempf`; Without this assignment to `dd, params->dangle5[i][j]` will have different values. Depending upon the compiler, the new value may perhaps be similar to those in `dangle3`.

#### 3.3.3 <_params.c_339>

This causes line 339 in `params.c` to be deleted. Without `params->mismatchExt[i][j][k] =(mm > 0) ? 0: mm`; array `mismatchExt` may be left at its initial values (0).

#### 3.3.4 <_model.c_860>

This mutation deletes line 860 from file `model.c` (i.e. `md->noLP = noLonelyPairs;`). This means model parameter `noLP` is not set and instead retains its default value (0).

### 3.4 Lesson

All four code changes found by evolution work on the quality of RNAfold’s predictions indirectly. They each modify the values of the energy parameters used in the dynamic programming’s energy calculation. In the next set of experiments [29], instead of mutating the C code itself, we ask GGGP to directly evolve the more than 50,000 parameters in RNAfold’s model of RNA.

## Acknowledgements

I am grateful for the assistance of John Andrews, Neil Daeche, Rhiju Das Fernando Portela and Ronny Lorenz.

1 To use the SSE code, ViennaRNA must be configured with ./`configure --enable-sse` when it is compiled. https://www.tbi.univie.ac.at/RNA/documentation.html

